# Designer Scaffolds for Interfacial Bioengineering

**DOI:** 10.1101/2020.11.06.371278

**Authors:** Ryan J. Hickey, Maxime Leblanc Latour, James L. Harden, Andrew E. Pelling

## Abstract

In regenerative medicine, the healing of the interfacial zone between tissues is a major challenge, yet approaches for studying the complex microenvironment of this interface remain lacking. Here, we study these complex living interfaces by manufacturing modular “blocks” of naturally porous decellularized plant-derived scaffolds with a computer numerical controlled mill. We demonstrate how each scaffold can be seeded with different cell types and easily assembled in a manner akin to LEGO™ bricks to create an engineered tissue interface (ETI). Cells migrate across the interface formed between an empty scaffold and a scaffold pre-seeded with cells. However, when both scaffolds contain cells, only a shallow cross-over zone of cell infiltration forms at the interface. As a proof-of-concept study, we use ETIs to investigate the interaction between lab grown bone and connective tissues. Consistent with the above, a cross-over zone of the two distinct cell types forms at the interface between scaffolds, otherwise the populations remain distinct. Finally, we demonstrate how ETIs are biocompatible in vivo, becoming vascularized and integrated into surrounding tissue after implantation. This work creates new tissue design avenues for understanding biological processes or the development of synthetic artificial tissues.

## 1. Introduction

The body consists of a variety of tissues, and the interface between them presents a unique juxtaposition of interacting cellular environments^[1–3]^. Traditionally, bioengineers design scaffolds for one environment, however there is also a need to develop model systems to create and investigate tissue interfaces^[4–8]^. Recently, greater attention to the interfacial zone between tissues has attracted more attention. Hydrogel interfacing mechanisms along with innovative surface chemistries and covalent bonding mechanisms have been explored to address the hydrogel-tissue interface issues preventing restoration of tissues to intact states after damage^[9]^.

Highly organized and graded tissue interfaces prevent abrupt transitions between different tissues and modulate stress concentration. Natural interfaces are characterized by changes in local microarchitectural features and compositions which play an important role in governing the interaction between two distinct tissues. Additive manufacturing (AM) strategies to implement these design features, such as layering, 3D printing, fluid mixing and electrospinning, have been used to create biomimetic tissue interfaces^[10,11]^. These strategies, each with their own set of advantages and disadvantages, have rapidly progressed the field of tissue interface engineering due to their versatility and high spatiotemporal resolution. Beyond additive manufacturing approaches, other strategies to create gradients for tissue interface construction have been investigated as well^[10,11,20,21,12–19]^.

The modular LEGO™-like assembly of materials and hydrogels for tissue engineering has been previously explored in the literature at a variety of length scales. Specifically, LEGO™-like multilayered artificial tissues have been created with photon induced polymerization and miniaturized modular microcage scaffolds produced with lithography-based 3D printing^[22–24]^. The stacking of distinct modules allows for numerous configurations and scales of materials to be fabricated. Inspired by this, we constructed living engineered tissue interfaces (ETIs) by combining highly porous, decellularized, plant-derived scaffolds in a modular fashion. Plant-derived scaffolds can be selected to mimic natural human structures in vitro and in vivo^[25,26,35,36,27–34]^. Intense research has been focused on designing biomaterials to enhance tissue integration in specific cases^[37–40]^; however, the widespread potential of this technology necessitates a platform that can be adapted to suit particular biomedical needs. We hypothesized that the modular assembly of plant-derived biomaterial subunits could create effective ETIs.

The physical environment affects many cellular processes including differentiation, migration, force generation, proliferation, spreading, alignment, and gene expression^[41–44]^. The interplay between the physical characteristics of the microenvironment and biochemical signaling is complex, and extensive research has been dedicated to elucidating the mechanisms responsible for these phenomena^[45–49]^. These insights have highlighted a need to develop strategies for the fabrication of biomimetic constructs that more closely match the mechanical properties and microstructure of the native environment of the cell^[50]^.

Another layer of complexity is added when considering the physical interface between tissues. These regions often bridge at least two biophysically distinct environments, and the transitional zone has its own multifaceted organization. For example, tissue entheses contain graded insertion sites with uncalcified and calcified regions, different extracellular matrix proteins (e.g. collagen type), and specific fibrillar arrangements in order to create a graduated transition between stiff and soft tissues; these interfacial zones are susceptible to injury and are difficult to repair and regenerate^[1,51–53]^. Although damage to the interfaces between distinct tissues and structures within the body can have significant consequences, there are few, if any, bioengineering strategies to study their regeneration from the perspective of basic fundamental science or clinical translation. Therefore, development of novel strategies to biofabricate artificial tissue interfaces may lend novel insights into how such regions may be designed, engineered, and repaired in a regenerative medicine context.

Moreover, when attempting to create designer biomaterials and scaffolds for fabricating tissue interfaces it is important to also consider the mechanical performance and matching of physical properties with the surrounding environment. Such considerations are integral for effective biomaterial design and performance^[54–56]^ as mechanical/physical mismatch can lead to poor tissue integration which is ultimately a leading cause of implant failure^[52]^. Additionally, the transitional zones between tissues are exposed to a wide variety of mechanical forces^[57–60]^. Tools to study the responses to stresses and strains applied to these interfaces are required to advance our understanding of how these regions respond to compression, tension, and shear. Designer scaffolds for artificial tissue interfaces can help tease apart the gaps in knowledge surrounding tissue interfaces. Growing two or more distinct tissues and bringing them together into one functional unit as an ETI presents exciting possibilities to design artificial tissues that have analogues in the human body, but going a step further, engineering tissues that have unique physical properties.

Our previous work reveals that the decellularized plant-derived scaffolds have a Young’s modulus that is comparable to that of various human tissues and cell types; moreover, specific plant candidates can be chosen to mimic particular structures in the body. Nevertheless, simple compression measurements fail to fully characterize the mechanical properties of these scaffolds. In the body, our tissues are subjected to complex shear forces in addition to simple tensile and compressive loads. Here, we provide a more thorough mechanical characterization of the ETI. This platform for ETI design establishes a “plug and play” method for combining multiple cell types and distinct physical environments.

## 2. Results

### 2.1. Engineered Tissue Interface (ETI) Fabrication

In order to create ETIs, we combined subunit components without the use of additional crosslinkers or glues. Interlocking composite pieces were fabricated to yield scaffolds that could be repopulated with distinct or identical cell types and subsequently re-combined into a single unit (**Figure 1**)^[27]^. To create such scaffolds, we employed 3D milling with a computer numerical controlled (CNC) router to reproducibly carve the structures from apple hypanthium tissue (Figure 1a, Video S1). A simple design for interlocking materials is the common stud and anti-stud geometry of LEGO™ bricks (Figure 1b and **Figure S1**). We designed single stud and anti-stud bricks, wherein the stud subunit consisted of a 5 mm x 5 mm x 2 mm base with a 2 mm (height and diameter) cylindrical peg protruding from the center, and the anti-stud subunit had a 2 mm diameter cylindrical hole in the center of the 5 mm x 5 mm x 2 mm base (Figure 1b). Tissue interface formation was a direct consequence of interlocking the components together (Figure 1c). Prior to assembly, the individual subunits were decellularized, processed, and repopulated with mammalian cells as previously described^[27]^ (Figure 1d and **Figure S2**).

**Figure 1:**
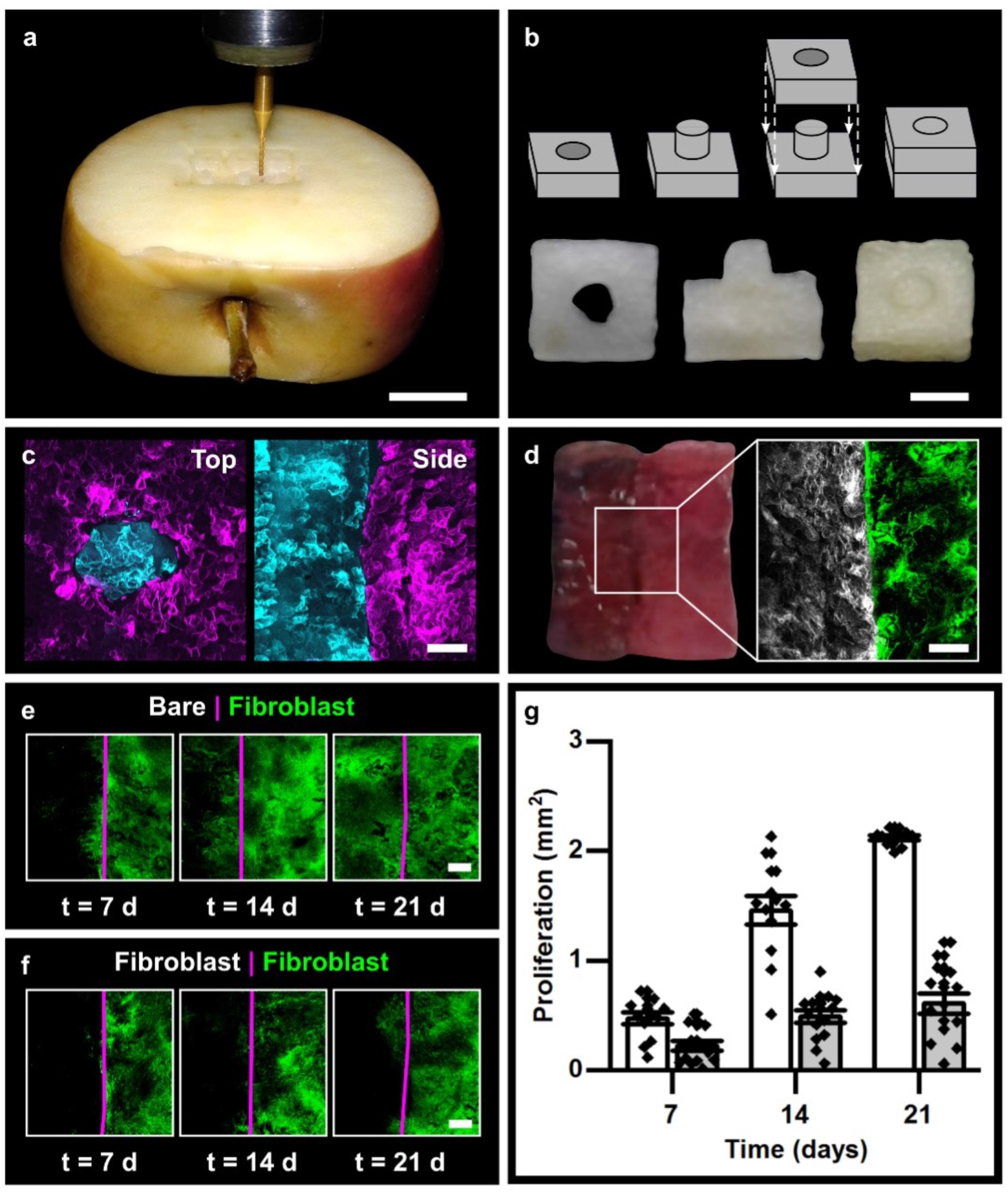
Interlocked biomaterials for ETI formation. **(a)** A CNC router was used to carve the interlocking materials. Scale = 15 mm. **(b)** Geometry of the single stud and anti-stud interlocked units as well as the decellularized subunits and the combined entity. Scale = 2.5 mm. **(c)** Maximum z-projected confocal image of the top view and side view of the interlocked unit with one subunit stained with Calcofluor white (cyan) and the other stained with Congo Red (magenta), scale = 500 µm. **(d)** Tissue interface formation: left = side view of interlocked unit, right = maximum z-projected confocal image of the two cell populations. White = Hoechst 33342 stained nuclei of NIH 3T3 cells, green = GFP NIH 3T3 cells. Scale = 500 µm. Fibroblast cell (green) migration across the interface (magenta) of the interlocked unit with **(e)** one subunit preloaded with cells and the other void of cells, and **(f)** both loaded with cells (only one population transfected with GFP). Scale = 500 µm. **(g)** Quantification of cell migration across the interface. White = bare | fibroblast, grey = fibroblast | fibroblast, N = 12.

### 2.2. Cell Migration

Cells must be able to traverse the interface and interact with the adjacent population when required^[61]^. Controlled in vitro cell migration assays were carried out, which revealed that when a scaffold pre-loaded with fibroblast cells was interlocked with a bare subunit, the cells migrated across the interface and invaded the bare scaffold (Figure 1e). The proliferation was monitored weekly for three weeks, and each week presented a significant increase in cell infiltration (p < 0.001) (Figure 1e, g). After three weeks of culture, the initially barren scaffold had a comparable cell number density to the pre-seeded subunit (Figure 1e, g and **Figure S3**). In order to assess the uniformity of the cell migration across the scaffolds, 200 µm wide regions of interest (ROIs) from the edge of the interface were evaluated (**Figure S3)**. The total cell area coverage across each of the ROIs was used to normalize the data and calculate a relative percentage of cells in each ROI. The relatively large initial variance at 7 days increased by 14 days, and then it significantly decreased by 21 days when near full coverage was achieved.

We repeated the cell migration assays with fibroblasts loaded on both subunits. The two fibroblast groups were distinguishable, as only one population contained green fluorescent protein (GFP) (Figure 1d-f). We found that GFP-labelled cells crossed the interface; however, when comparing the dual and single cell migration assays, there was significantly less migration when both scaffolds were pre-loaded with cells (p < 0.002), in which case the cells did not fully invade the adjacent scaffold. Therefore, the two cell populations remained largely distinct away from the interface on their respective sides but were well integrated at the interface (Figure 1e-g).

### 2.3. Bone-connective tissue interface

ETIs can be designed to replicate specific tissue interfaces, such as the bone-connective tissue interface, which delineates two separate regimes (**Figure 2**). This was achieved by culturing MC 3T3 E1 subclone 4 pre-osteoblasts on one subunit to confluency followed by differentiation into osteoblasts, which mineralized the scaffold. An opposing scaffold was also cultured until confluence with mouse NIH 3T3 fibroblasts and interlocked with the calcified subunit (Figure 2). As osteoblasts deposit calcium and mineralize their extracellular matrix, there are different biochemical and physical environments of the two cell populations forming the bone-connective tissue ETI. Alkaline phosphatase and calcium staining revealed that the composite was comprised of two distinct regions away from the interface (Figure 2 and **Figure S4**). We then probed the local cellular environments using scanning electron microscopy (SEM) and electron dispersion spectroscopy (EDS). SEM revealed two distinct surface topographies (Figure 2c, f and **Figure S5**). On the bone component, a smooth coating was deposited on the scaffold (Figure 2c, f and Figure S5). This coating was absent on the fibroblast side; instead, thick layers of cells were observed (Figure 2c, f and Figure S5). EDS was employed to confirm that the smooth coating on the differentiated bone component was mineralization. The characteristic X-rays confirmed the presence of calcium (∼3.69 keV) and phosphorous (∼2.01 keV) on the bone scaffolds (Figure 2g). These elements were not present to a detectable level on the fibroblast side (Figure 2h, i). The interfacial region was also devoid of mineralization.

**Figure 2:**
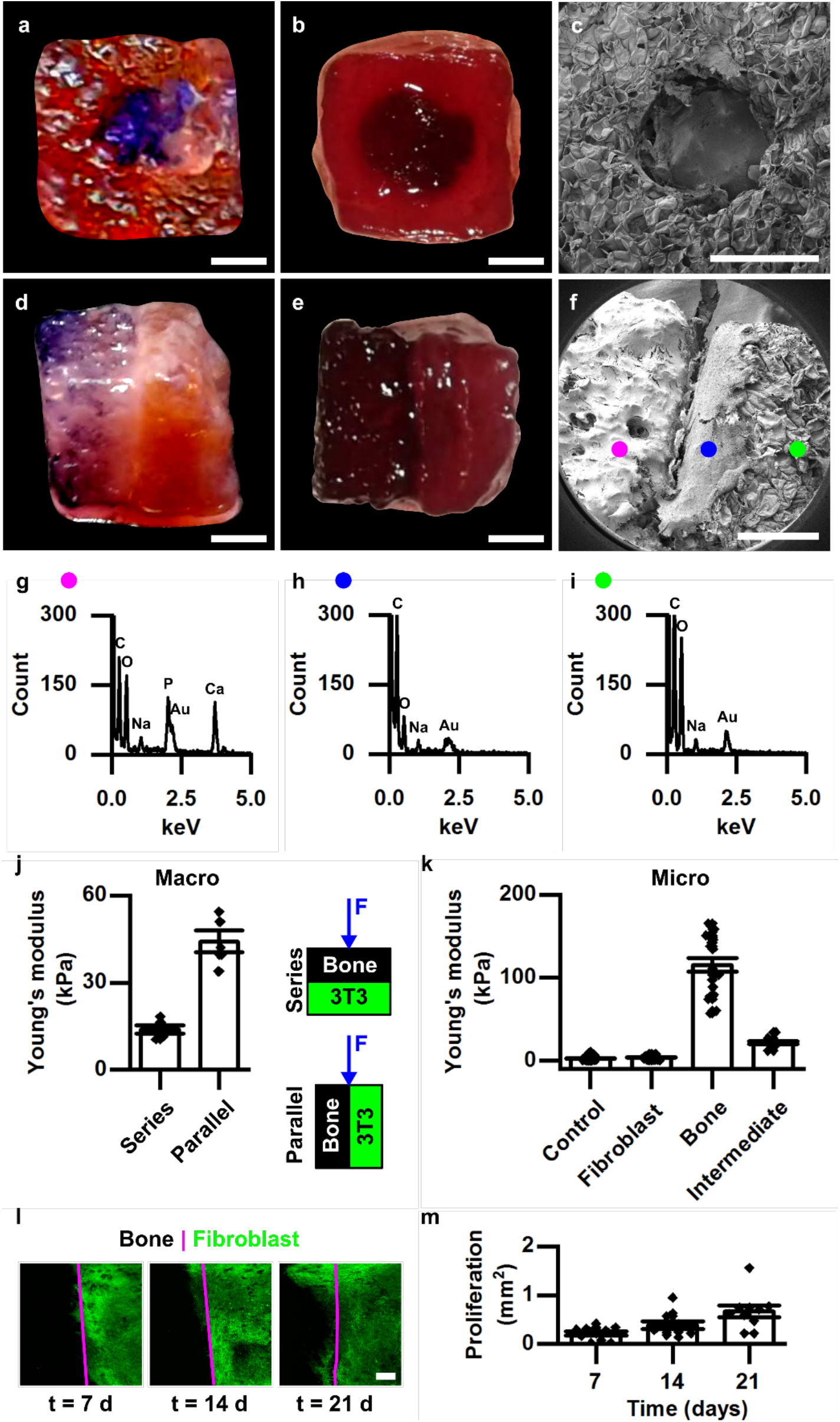
Bone-connective tissue ETI. **(a)** Top view of scaffolds stained to reveal alkaline phosphate. Purple = positive for alkaline phosphatase activity, orange/red = negative. **(b)** Top view of scaffolds stained for the presence of calcium staining. Deep red = calcified scaffold, light red = non-calcified. **(c)** Top view of SEM of bone-connective tissue ETI with smooth mineralized peg inserted into the non-mineralized fibroblast covered subunit. Corresponding side views of the scaffolds stained for **(d)** alkaline phosphatase and **(e)** calcium. **(f)** Corresponding side view of the scaffold imaged with. Scale bar **(a-f)** = 1 mm. **(g-i)** EDS chemical analysis of the corresponding regions (magenta = bone subunit, blue = fibroblast covered subunit adjacent to the interface, green = fibroblast covered subunit). **(j)** Compressive moduli of the bone-fibroblast ETI compressed in the series and parallel directions, N = 5. **(k)** Local mechanical probing: nanoindentations with AFM. **(l)** Fibroblast cell (GFP NIH 3T3) migration across the bone-connective tissue ETI, scale = 500 µm. **(m)** Quantification of cell migration, N = 10.

The separation of the mineralized and non-mineralized tissue resulted in two regions with different mechanical profiles (Figure 2). The composite is composed of materials with different elastic moduli; therefore, an effective Young’s modulus can be obtained from the geometry, direction of applied force, and the moduli of the constituents (Figure 2j). To illustrate this, force was applied normal to the interface (series orientation) or parallel to it. The effective Young’s modulus in the series arrangement was 13.8 ± 3.3 kPa which was significantly less than the parallel conformation with a value of 44.2 ± 8.4 (p < 0.001). ETIs consisting of components with different mechanical profiles allows stress shielding biomaterials to be formed^[1,54,55]^ (Figure 2j and **Figure S6**). In addition to macroscale compression, atomic force microscopy was used to probe the local mechanical properties in both regions (Figure 2k); the two stiffness regimes observed were comparable to the reported literature values for bone and fibroblast cells ^[62,63]^. The bone component had a Young’s modulus of 115.0 ± 8.2 kPa, whereas the fibroblast component had a significantly lower modulus of 3.9 ± 0.3 kPa (p < 0.001). Interestingly, there was an intermediate stiffness found in the interfacial region with a modulus of 21.7 ± 1.7 kPa, as expected for a region composed of fibroblasts, undifferentiated pre-osteoblasts, and differentiated osteoblasts surrounded by their mineralized matrix. A cell migration assay from the fibroblast component onto the bone section demonstrated that cells intermixed at the interface and exhibited a 13% proliferation increase as in the previous experiment (p = 0.971, compared to the dual fibroblast migration assay), which is similar to the natural bone-connective tissue interface^[64,65]^ (Figure 2l, m).

### 2.4. Mechanical characterization of ETIs

Mechanical characterization is integral for designing tissue interfaces with specialized structure-function relationships^[65]^. Three separate mechanical characterizations are of particular interest for these interlocked composite materials: compression, tension, and shear (**Figure 3a-f**). We examined scaffolds with and without added fibroblast cells in the blocks. Compression tests probe the bulk elastic properties of the two blocks in series, and the interface between blocks plays a minor role. It was found that the presence of cells had no significant impact on the apparent stiffness of the material: 24.4 ± 1.3 kPa, (p = 0.68) (Figure 3g), as intrinsic cell elasticity is relatively weak compared to that of the matrix material. Under tension, the mechanical response is dominated by the cohesiveness of the interfacial region. This can be seen clearly in Figure 3e, which shows a progressively decreasing contact area between blocks as two blocks separate under the applied tension. In this case, the stress required to initiate separation of the two subunits was greater when fibroblasts were seeded on the scaffolds (cells: 1.56 ± 0.1 kPa, control: 0.6 ± 0.1 kPa, p < 0.001) (Figure 3h), as cell-matrix and cell-cell adhesion both contribute to the mechanical integrity of the interfacial region. Applied shear provides a direct probe of interfacial cohesion and integrity. Controlled steady shear rate experiments were characterized by an initial reversible response, where stress σ is found to be a sub-linear power law of applied strain ε, σ∼ε^y^ with y β 3/4, followed by subsequent strain softening (Figure 3i). The lack of a linear stress-strain relationship at low strains is indicative of mechanics dominated by stick-slip events at the interface. For larger strains, eventually transient failure in the slip plane of the interface occurs, which is followed by a steady state friction response (**Figure S7**). Interestingly, the power-law stress-strain response was found to be reversible even after failure in the slip plane for both relatively slow and fast applied strain rates (Figure S7), which may serve as a valuable design feature for preventing the scaffolds from irreversible fracturing under large loads in situ. In shear conditions, the presence of fibroblast cells did not qualitatively change the stress-strain relationship; however, samples with cells were found to be slightly more compliant (Figure 3i). An interfacial region containing cells, which have a relatively lower intrinsic modulus, may be more compliant in shear deformation than an interface composed of pure matrix.

**Figure 3.**
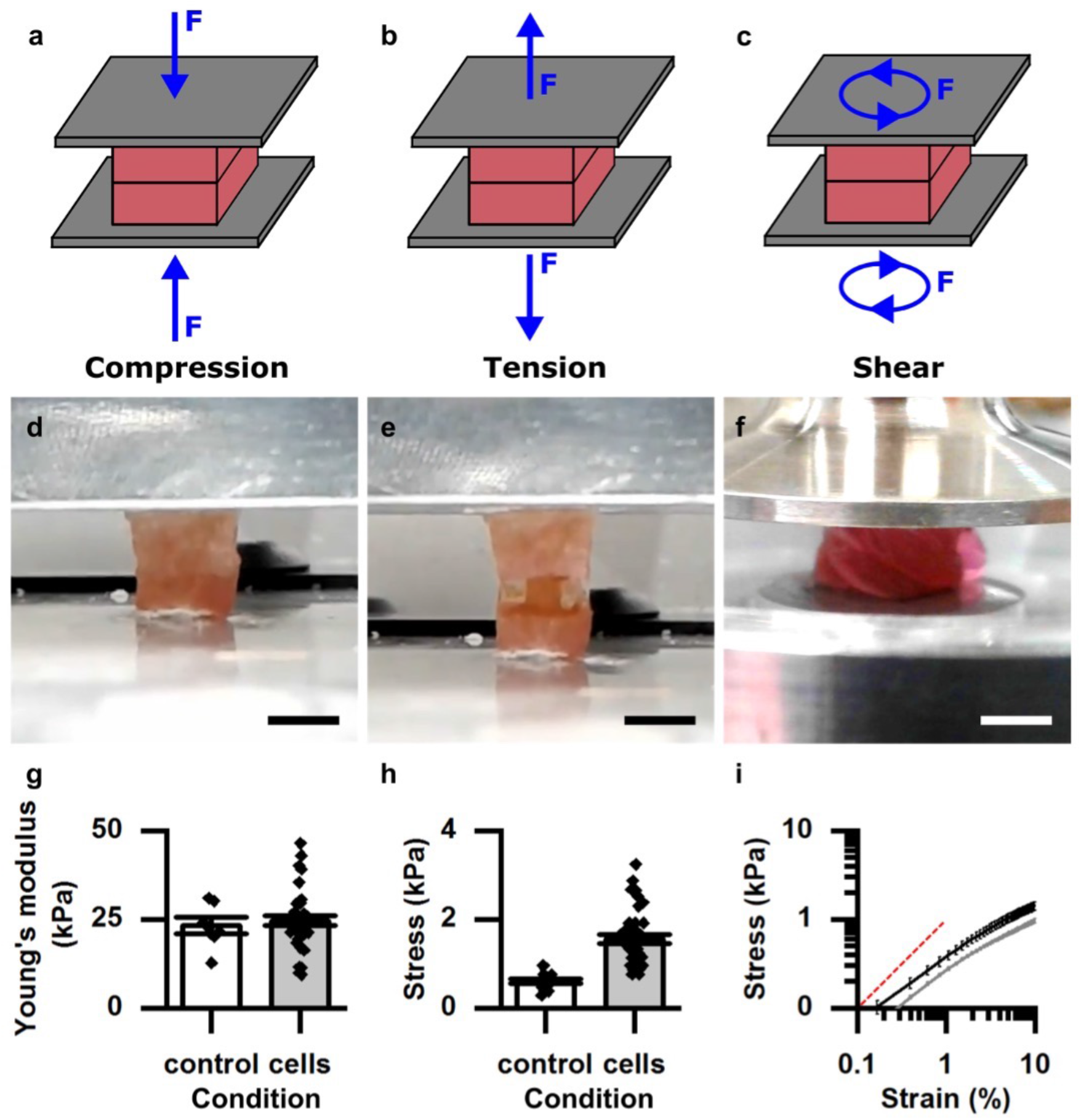
Mechanical testing of ETIs. **(a-c)** Cartoon illustrations of the three classes of mechanical tests that were performed: compression, tension, and shear. **(d-f)** Interlocked ETIs under compressive, tensile, and shear stresses. Scale = 4 mm. **(g)** Compressive Young’s modulus with (N = 36) and without fibroblast cells (N = 7) with cells. **(h)** Maximum tensile stress prior to subunit separation with (N = 39) and without fibroblast cells (N = 15). **(i)** Shear stress σ vs strain ε in a controlled steady shear rate experiment at a strain rate of 0.01 s^-1^, N = 3. Black = without cells, grey = with cells. The red dashed line is provided as a guide for the linear regime. Power-law behavior is observed at low strains: σ∼ε^y^ for y = 0.74 ± 0.02 without cells and y = 0.72 ± 0.02 with cells.

### 2.5. In vivo implantation

The potential applications of ETIs are not limited to *in vitro* studies of tissue interfaces; we foresee ETIs playing an essential role in biomaterial implant design. As a proof-of-concept experiment, we implanted the interlocked composite materials subcutaneously in an immunocompetent rat model for 4 weeks^[26,27]^ (**Figure 4a**). The subunits remained interlocked despite being placed under the skin (Figure 4b). Histological analysis revealed that the cells invaded the peripheral regions of the biomaterials, and blood vessel formation was clearly observed (Figure 4c-f and **Figure S8**)^[26,27]^. The invading fibroblasts from the surrounding animal tissue laid down their own collagen network within the cellulose-based scaffold (Figure 4d, f).

**Figure 4.**
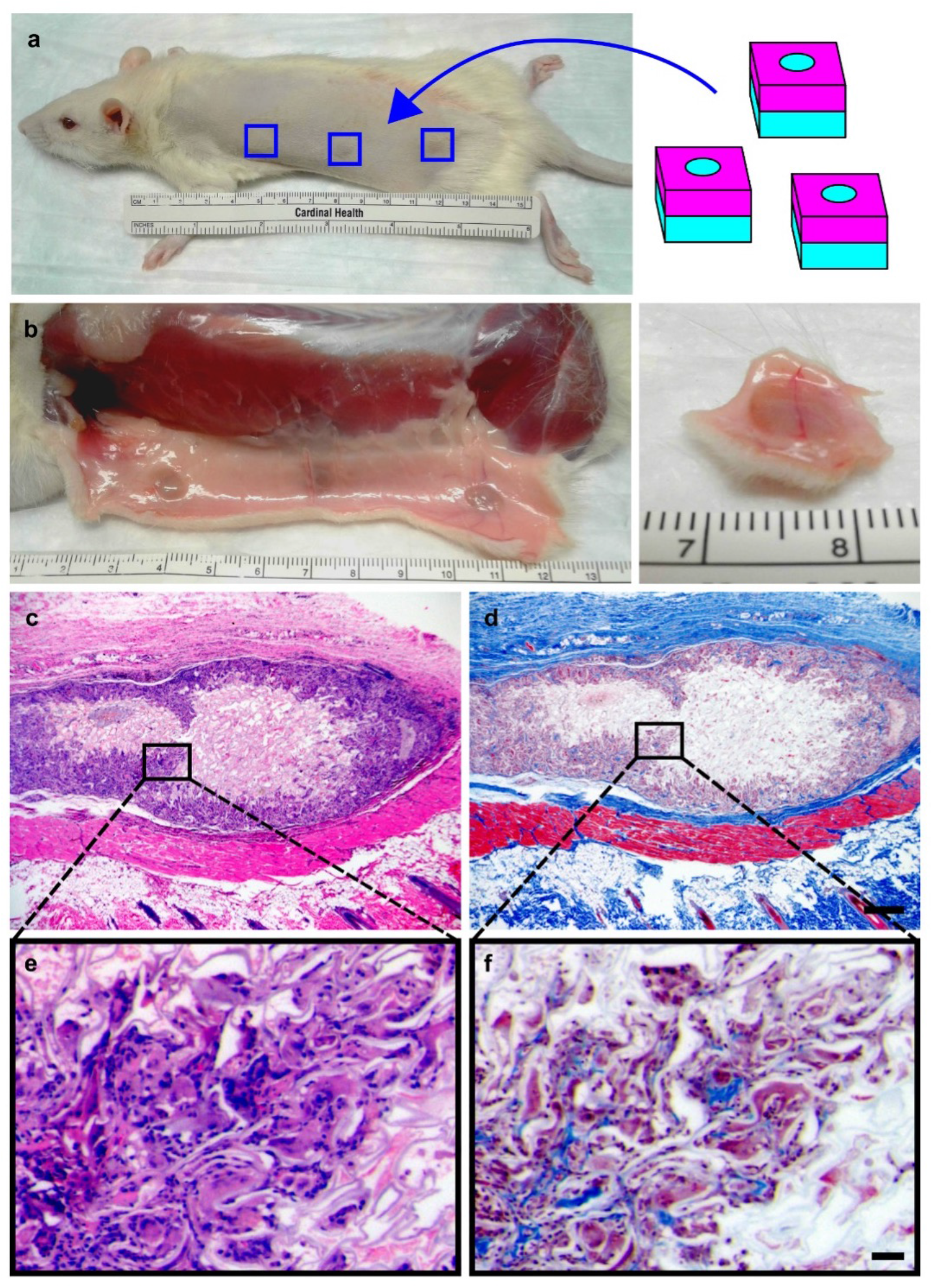
In vivo implantation of ETIs. **(a)** Schematic showing the location of three subcutaneous implanted ETIs in a rat model. **(b)** ETI sample resection showing the subunits remained interlocked and underwent compression after 4 weeks. **(c, e)** H&E staining revealing substantial of the cell invasion in the outer regions of the interlocked biomaterial. **(d, f)** Masson’s Trichrome staining showing collagen deposition (blue) on the scaffold. Scale **(c, d)** = 500 µm, **(e, f)** = 50 µm.

## 3. Discussion

As demonstrated in this study, plant-derived ETIs establish a novel approach for studying and implementing tissue interface models. The main advance in this particular study is that the interlocking biomaterials did not need to be 3D printed or fabricated utilizing complex techniques in order to create porosity and interlocking abilities. Rather, plant-derived scaffolds are naturally porous, allowing for rapid prototyping of macroscale interlocking “blocks” which allow for cell loading, infiltration, and proliferation in a highly customizable manner. In addition, decellularized plant-based biomaterials are highly accessible and simple to produce. The approaches presented here allow many others to now employ simply produced, naturally porous, interlocking biomaterials in future studies of the design and engineering of artificial biomimetic tissues.

Significant insights in matrix composition, mechanical loading, and cell organization have been gained from specific tissue interface models of entheses, such as the lack of transition and insertion zones at interfaces post-operation^[1,37,66,38–40,51–53,61,65]^, yet platforms to study tissue interfaces are lacking. The choice of the LEGO style used design here was to demonstrate proof-of-concept, but one can generate specific interlocking geometries to suit particular applications (**Figure S9**). Although the LEGO brick is artificial and irregular on a macroscopic scale, it provides a simple demonstration of creating interfaces between subunits. By linking together distinct block types, engineered interface construction was made possible. The use of plant-derived, porous, cellulose-based scaffolds results in interesting geometries, specifically at the interface. Importantly, on a microscopic scale, the irregular shape of the blocks becomes irrelevant and arbitrary, as the interconnected porous cellulose structure creates a 3D scaffold to support cell growth within and across the interface between interlocked subunits. Several groups have now demonstrated that that existing plant structures can be selected and repurposed to nearly match the geometry of animal tissues^[25,26,35,36,27–34]^. Interlocking different subunits with different microstructures to create designer interfaces increases the potential applications of these materials. As scale is a challenge in biomaterial design, this approach circumvents the need to source large starting materials or to decellularize large tissues.

We have shown that decellularized plant-derived scaffolds can be shaped into complementary interlocking subunits, and that cells can migrate across the interface between such subunits. A combination of local geometrical factors of the interface and leading cell effects of cell population migration are plausible causes of the non-uniform “wavefront” of advancement observed (**Figure S3**). In the absence of an existing cell population, such as an implanted biomaterial void of cells, this extensive cell migration and infiltration is usually desirable^[67]^. When a neighbouring cell population was present, migration across the interface was observed, yet the migrating cells did not fully invade the adjacent scaffold. This result is pertinent for tissue interfaces, as a uniform co-culture of cells does not replicate entheses in vivo. Two obstacles in entheses design are the achievement of adequate cell invasion and the creation of an appropriate transitional zone between segregated microenvironments^[68–70]^. The modular assembly with porous scaffolds enables cells from two distinct regions to form an intermediate zone at the interface between the subunits^[66]^. The use of these ETIs has significant implications for future biomaterial design including the ability to fabricate cellular organization that is not easily achieved with conventional mixed cell culture techniques.

In this study we utilized ETIs to study the specific case of a bone-connective tissue interface. Bone cells are surrounded by a mineralized extracellular network; however, this mineralization is absent in the connective tissue^[37–40,51–53]^. A lack of integration of the two regions is a key factor in implant failure^[52]^. We showed that cells migrated across the interface and that distinct mechanical environments were established. In this interfacial region between the two scaffolds, no significant mineralization was observed. In vivo, there is a similar zone in the enthesis known as the uncalcified avascular fibrocartilage zone, which is a non-mineralized, avascular region found between the fibrous connective tissue and the tidemark separating the hard and soft tissues^[64]^. Several models have been developed in an attempt to recreate this interface which consists of creating different environments within a single stratified material or two or more adjacent components^[37–40,51–53]^. Nevertheless, a universal “plug and play” system has not been established for engineering and designing models of such tissue interfaces.

Spatial organization and transitional zones within tissues and tissue interfaces are highly complex and integral to proper function; thus, the ability to combine them in a spatially controlled manner is essential. The modular assembly of components into a functional unit allows customizable designer interfaces to be formed. The choice of scaffold base material and interlocking format are important considerations when developing the artificial tissue enthesis. Here, we observed that the presence of cells increased the integration of the two subunits, as evidenced by tension experiments; however, the presence of cells did not significantly change the results of the compression and rotation tests. Of course, such mechanical properties are likely to be highly dependent on the specific scenario being investigated, especially when different cell types and/or interlocking geometries (e.g. multi-post blocks) are being investigated.

Importantly, we also demonstrate that the ETIs presented here are suitable for in vivo applications. It was found that the interlocked material did not disassemble after implantation in an animal model. These results allow for speculation about how these materials may be used to replace or regenerate tissue entheses in the body; however, more studies will obviously be required. From a different angle, these materials also offer the ability to study complex systems and processes associated with tissue interface barriers, which can be found throughout the body, such as in the kidney, liver, intestines, and the brain. Significant advances have been made in microfluidics to study these tissue barriers; however, the focus is almost entirely on the fluid-barrier interface. By using ETIs in a combinatorial approach with the recent microfluidic advances, a deeper understanding of how flow across the barrier can affect cells of stratified tissues may be achieved^[71–73]^. The modular assembly approach of ETIs presented here ultimately provides an exciting new framework for the design of living tissues.

Here we demonstrated the modular assembly of premade solid units as a novel alternative to 3D printing from the liquid state. That being said, 3D bioprinting is a useful approach to designing multicompartment scaffolds for interface tissue engineering as well. Both approaches have advantages and disadvantages, yet the most exciting prospect that arises from developing these techniques is to use them in combination to exploit the advantages of both methods simultaneously when creating designer artificial tissues for numerous potential applications.

## 4. Experimental Section/Methods

### Scaffold production

A Shapeoko 3 CNC router with a 0.8 mm, 180° drill bit was used to carve McIntosh Red apples (Canada Fancy) into arrays of complementary stud and anti-stud geometries. Cutting was performed at a speed of 1 mm/s. A Mandolin slicer was then used to cut the arrays of the subunits to their desired thickness. The stud piece consisted of a 5 mm x 5 mm x 2mm base with a 2 mm cylindrical peg protruding from the center, and the anti-stud subunit had a 2 mm cylindrical hole in the center of the 5 mm x 5 mm x 2mm base. As done previously, the samples were transferred to a 0.1% SDS solution and decellularized for 48 h while being shaken at 180 RPM, then were washed three times with dH_2_O. Following the washes, the subunits were incubated in 100 mM CaCl_2_ for 24 h and were then washed three times with dH_2_O to remove the salt residue. To sterilize the samples, a 70% ethanol incubation was performed, followed by three more wash cycles with dH_2_O.

### Cell culture

All cells were maintained at 37°C and 5% CO_2_. NIH 3T3 fibroblast cells (ATCC NIH/3T3 CRL-1658, from *Mus musculus*) were cultured in Dulbecco’s Modified Eagle Medium – High Glucose (DMEM), supplemented with 10% fetal bovine serum and 1% penicillin/streptomycin (100 U/mL and 100 μg/mL respectively) (Hyclone Laboratories Inc.). Conversely, the MC 3T3 E1 Subclone 4 pre-osteoblast cells (ATCC CRL-2593, from *Mus musculus*) were culture in MEM-alpha supplemented with 10% fetal bovine serum and 1% penicillin/streptomycin (100 U/mL and 100 μg/mL respectively). In order to invoke differentiation of the pre-osteoblasts, 4 mM inorganic phosphate (Sigma) and 50 μg/mL ascorbic acid (Sigma) were added. For sub-culturing, cells cultured on cell culture plates were trypsinized and resuspended in the appropriate medium. The cells were counted and centrifuged in order to separate the cells from the trypsin and the media. The supernatant was aspirated, and the pellet containing 5×10^4^ cells was resuspended in fresh culture medium. The cells were seeded onto the biomaterial, and the cells were allowed to proliferate and invade the scaffold for 2 weeks prior to interlocking the complementary subunits. After 2 weeks, the subunits were manually clicked together using sterile tweezers by pressing down the stud subunit into the hole of the anti-stud piece. The culture media was replaced every day and the samples were transferred to new culture plates after 1 week of growth.

### Confocal microscopy

The biomaterials were imaged with a Nikon TiE A1-R confocal microscope. The plant-derived scaffold was visualized with Calcofluor white staining (30 min, 1 µg/mL, Sigma) and Congo Red (30 min, 0.1 µg/mL, Sigma). Cell nuclei were stained with Hoechst 33342 (Invitrogen) (5 min incubation, 10 µg/mL). ImageJ (Fiji) was used to process the images; brightness/contrast settings were adjusted to maximize the fluorophore signal.

### Cell migration assays

The confocal images of the GFP 3T3 cells were thresholded using the ImageJ (Fiji) adaptive threshold plugin, and the analyze particles plugin was used to measure the proliferation area on the adjacent scaffold. The data was normalized to the average scaffold area. The values presented are mean values ± the standard error of the mean (s.e.m.).

### Alkaline phosphatase staining

Prior to fixation, the scaffolds were washed with PBS. They were then fixed for 90 s with 3.5 % paraformaldehyde and then washed with wash buffer (0.05% Tween in PBS). The BCIP-NBT SigmaFast™) tablets were dissolved in dH_2_O and staining and imaging were completed within 1 h. During the staining, the samples were kept in the dark and were monitored. Once the staining was complete, the samples were washed and photographed.

### Alizarin Red S staining

Alizarin Red S (Sigma) was prepared by adding 1 g of the powder to 45 mL of dH_2_O. The pH was then adjusted to 4.3 with HCl and NaOH before raising the volume to 50 mL. Prior to staining, the samples were fixed as outlined above, except the duration of the fixation process was > 1h. The biomaterials were then washed with PBS. Calcium staining was performed with a 0.22 µm filtered Alizarin Red S stain pH 4.3 at a concentration of 200 mg/mL. The samples were submerged in the stain and incubated for 45 min. Following the calcium staining, the samples were thoroughly washed with dH_2_O until the colour ceased to run out of the samples. The samples imaged shortly afterwards.

### Scanning Electron Microscopy

Samples were fixed with 3.5 % paraformaldehyde for 48 h and washed with PBS. The samples were serially dehydrated with ethanol as indicated in Intech Open’s sample preparation guide^[74]^. The samples were dried with a samdri-PVT-3D critical point drier, and then gold-coated in a Hitachi E-1010 ion sputter. Scanning electron microscopy (SEM) and energy dispersive X-ray spectroscopy (EDS) were performed on a JOEL JSM-7500F field emission SEM at the Centre for Advanced Materials Research (CAMaR) at the University of Ottawa. The SEM images were recorded at 3 kV, and the EDS spectra were recorded at 15 kV.

### Compression and tension testing

The compressive Young’s modulus and the maximum tensile stress were measured using the compression and tension modes of a custom built dynamic mechanical analysis (DMA) device and LabVIEW. During the compression tests, the material was compressed to a 10% strain, at a strain rate of 50 μm/s. The force-displacement curves were converted to stress-strain curves, and they were fitted in Origin 8.5 to calculate the Young’s modulus. Likewise, the tensile measurements involved recording the minimum force required to separate the subunits at the same motor speed. In all DMA experiments, the samples were glued to the surface using Permabond Instant Adhesive 102 Medium Viscosity General Purpose Glue: a thin layer of the glue was evenly spread on the surface and the samples were gently pressed onto it and were incubated for 10 minutes prior to experimentation.

### Shear deformation testing

An MCR 301 Anton Paar Rheometer was used to determine the shear deformation behaviour of the materials. The samples were measured using a parallel-plate geometry with circular 12 mm plate tool diameters. For all shear experiments, the samples were glued to the surface using Permabond Instant Adhesive 102 Medium Viscosity General Purpose Glue: a thin layer of the glue was evenly spread on the surface and the samples were gently pressed onto it and were incubated for 10 minutes prior to experimentation. The samples were hydrated with CO_2_ independent media (25 mM HEPES) to prevent the samples from drying and resulting changes to scaffold mechanical properties. Shear stress was measured as a function of strain during a period of steady applied shear strain at a rate of 0.01 s^-1^.

### Atomic Force Microscopy

A Nanowizard II atomic force microscope (AFM) (JPK Instruments, Germany) was used to determine the substrate elasticity in all experiments. PNP-TR-50 cantilevers were used for each measurement and had an experimentally determined spring constant of 63.5 ± 7.2 mN/m. Force-indentation curves were acquired on substrates at 546 Hz with a set point of 1.0 nN. Substrate elasticity was calculated by fitting the force curves to the Sneddon-Hertz model for a conical indenter for shallow 200 nm indentations, assuming a Poisson ratio of 0.5 (PUNIAS 3D Software). For each substrate, N = 3 plates were prepared and 10 force curves were acquired at 25 random locations on the substrate for a total of 250 force curves.

### Animal surgeries and implantation

All procedures described in this study were approved and performed in accordance with standards, guidelines, and regulations set out by the University of Ottawa Animal Care and Veterinary Services ethical review committee. Results are also reported in accordance with ARRIVE guidelines. The subcutaneous implantation protocol was similar to what we have used previously. Briefly, immunocompetent rats were anesthetized using 2% Isoflurane USP-PPC (Pharmaceutical partners of Canada, Richmond, ON, Canada). ENDURE 400 Scrub-Stat4 Surgical Scrub (chlorhexidine gluconate, 4% solution; Ecolab Inc., Minnesota, USA) and Soluprep (2% w/v chlorhexidine and 70% v/v isopropyl alcohol; 3M Canada, London, ON, Canada) were used to prepare the shaved dorsal ventral area. Three 8 mm incisions were cut on the dorsal section of each rat, and a combined barren scaffold was placed in each incision. The incisions were then sutured using Surgipro II monofilament polypropylene 6–0 (Covidien, Massachusetts, USA). Transdermal bupivacaine 2% (as monohydrate; Chiron Compounding Pharmacy Inc., Guelph, ON, Canada) was topically applied to the surgery sites to prevent infection, and buprenorphine (0.03mg/ml; Chiron Compounding Pharmacy Inc. Guelph, ON, Canada) was administrated to alleviate pain. Animals were monitored and the sutures were removed after one week.

### Scaffold resections

At 4 weeks post-implantation, the mice were euthanized using CO_2_ inhalation and exsanguination via heart dissection. The dorsal skin was carefully resected and fixed in 10% formalin for at 72 h. The samples were then kept in 70% ethanol before being embedded in paraffin by the PALM Histology Core Facility of the University of Ottawa.

### Histology

Serial 4 μm thick microtome sections starting at the edge of the cellulose scaffolds were cut at 100 μm levels. The sections were stained with Hematoxylin and Eosin (H&E) and Masson’s Trichrome. For immunohistochemistry, the sections stained with Rabbit anti-CD45 (Abcam) or Rabbit anti-CD31 (Novus) were pre-treated using heat mediated antigen retrieval with a citrate buffer (pH 6.0, epitope retrieval solution 1) for 20 minutes. The sections were then incubated using a 1:1600 dilution (CD45) or 1:100 (CD31) for 30 minutes at room temperature and detected using an HRP conjugated compact polymer system. The slides were then stained with DAB as the chromogen, counterstained with Hematoxylin, mounted, and cover slipped.

### Statistical analysis

For multiple samples, one-way and two-way ANOVA tests were used to assess the statistical differences for samples with one or two factors respectively. The Tukey post hoc analysis was used to determine the value of the statistical difference between the individual samples. In the cases where only two sets of data were compared, the Student’s t-tests was used. All values presented are the mean ± the standard error of the mean (s.e.m.). Statistical significance refers to P < 0.05.

## Data availability

The research data required to reproduce these findings is available upon reasonable request from the corresponding author. The study is reported in accordance with ARRIVE guidelines.

## Supporting Information

Supporting Information is available from the Wiley Online Library or from the author.

## Acknowledgements

This work was supported by individual National Sciences and Engineering Research Council Discovery Grants awarded to AEP and JLH.

## Author contributions

RJH oversaw all experimental protocols and fabricated the scaffolds, performed cell culture, microscopy and analysis. MLL performed cell culture. MLL and RJH performed the surgical procedures. RJH and JLH performed rheological analyses. AEP conceived, directed and managed the project. All authors reviewed the data and prepared the manuscript for publication.

