## Supporting Information for "Designer Scaffolds for Interfacial Bioengineering"

### Supporting Information
### Designer Scaffolds for Interfacial Bioengineering

Ryan J. Hickey[1], Maxime Leblanc Latour[1], James L. Harden[1,2], and Andrew E. Pelling[1,3-5,*]

[1]*Department of Physics, STEM Complex, 150 Louis Pasteur Pvt, University of Ottawa, Ottawa, ON, K1N6N5 Canada*

[2]*Centre for Advanced Materials Research, University of Ottawa, Ottawa, ON, K1H8M5 Canada*

[3]*Department of Biology, Gendron Hall, 30 Marie Curie, University of Ottawa, Ottawa, ON, K1N5N5 Canada*

[4]*Institute for Science Society and Policy, Simard Hall, 60 University, University of Ottawa, Ottawa, ON, K1N5N5 Canada*

[5]*SymbioticA, School of Anatomy, Physiology and Human Biology, University of Western Australia, Perth, WA, 6009*

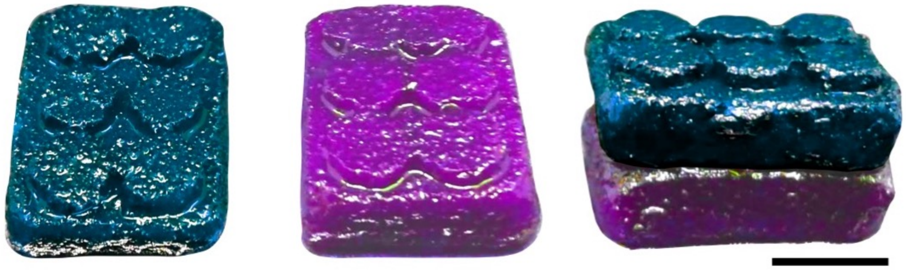

**Figure S1. Classic 3x2 Lego block design**. Two blocks were made from decellularized apple hypanthium tissue and were stained with Congo Red (magenta) and Calcofluor White (cyan). Scale = 1.5 cm.

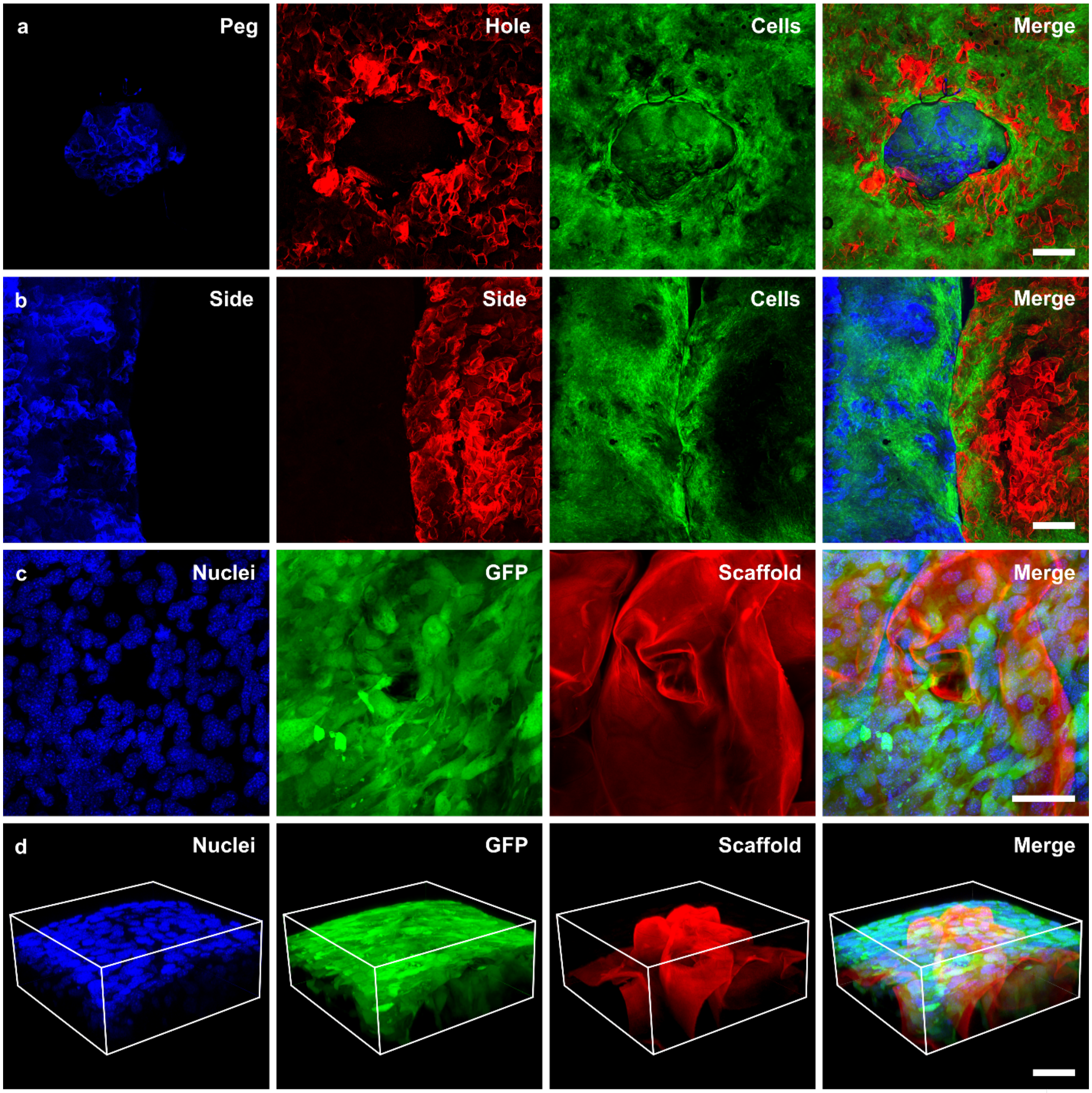


**Figure S2. Maximum intensity z-projected confocal images of interlocked ETIs and fibroblasts. (a)** Top view of interlocked peg (blue stained with Calcofluor white), hole (red stained with Congo Red), and GFP NIH 3T3 cells (green) on both subunits. **(b)** Side view of interlocked subunits. Subunit 1 (blue stained with Calcofluor white), subunit 2 (red stained with Congo Red), and GFP NIH 3T3 cells (green) on both subunits. Scale **(a,b)** = 500 µm. **(c)** Hoechst 33342 stained nuclei (blue), GFP NIH 3T3 cells (green), and scaffold stained with Congo Red (red). **(d)** 3D view of GFP NIH 3T3 cells on the cellulose-based scaffold. Blue = Hoechst 33342 stained nuclei, green = GFP, red = cellulose. Scale **(c,d)** = 50 µm.

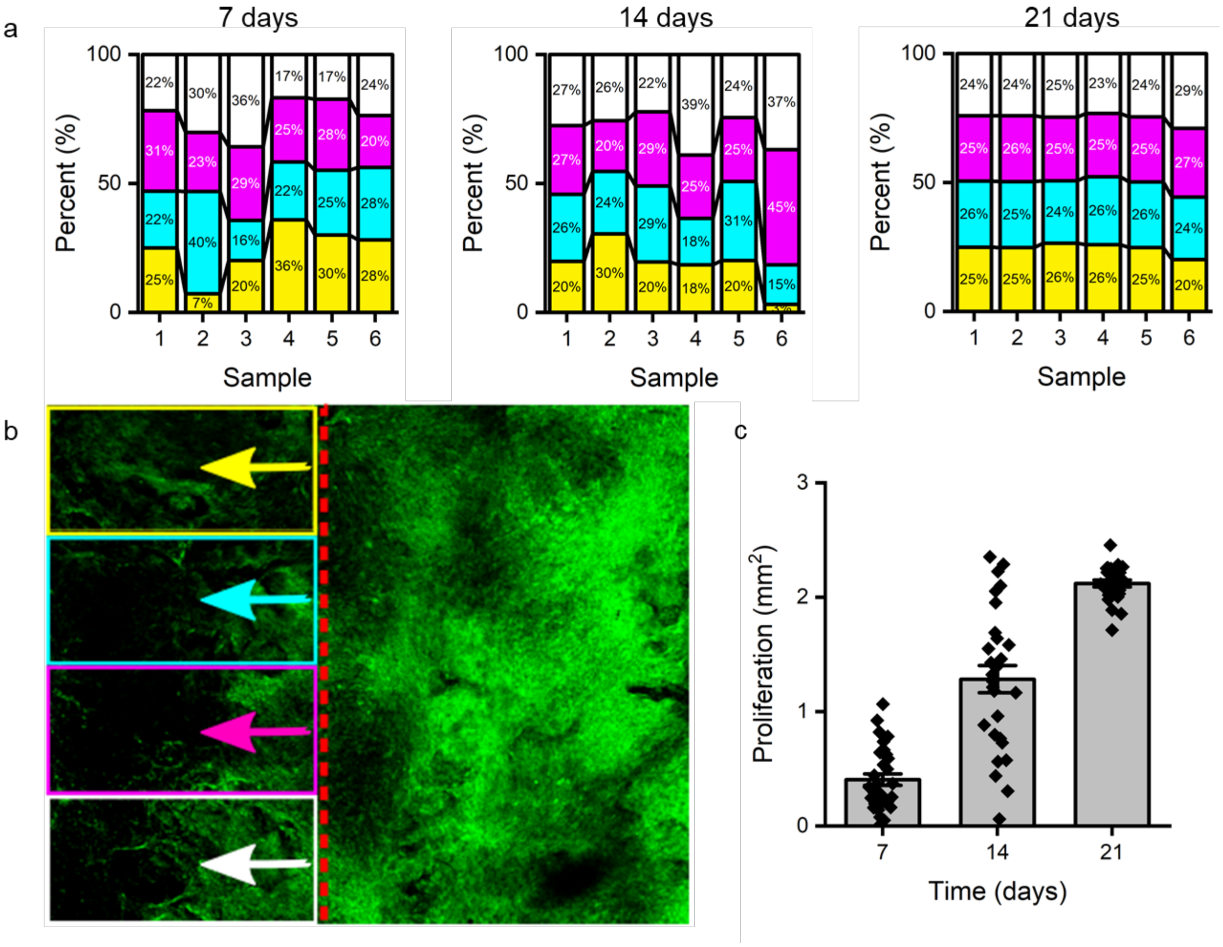


**Figure S3. Cell proliferation spatial variation.** The cell proliferation across the interface and onto the adjacent initially barren scaffold was evaluated by dividing the areas into 200 µm wide regions of interest (ROIs). This additional analysis gave insight into the uniformity of the cell migration across the interface. N = 6 samples were evaluated. **(a)** The stacked column plots depict the normalized relative cell coverage in each respective ROI (N = 4 ROIs). Each ROI is indicated by a different colour. **(b)** Visualization of the ROI selection on the initially barren scaffold. The ROIs are shown in different coloured rectangle boxes, and the principal direction of cell migration is indicated by the coloured arrows. The dashed red line marks the interface between the initially barren scaffold (left) and the pre-seeded scaffold (right). **(c)** Variance of the cell migration for different ROIs for the combined N = 6 dataset as a function of time. The columns of the scatter plot depict the mean and standard error of the mean (s.e.m.).

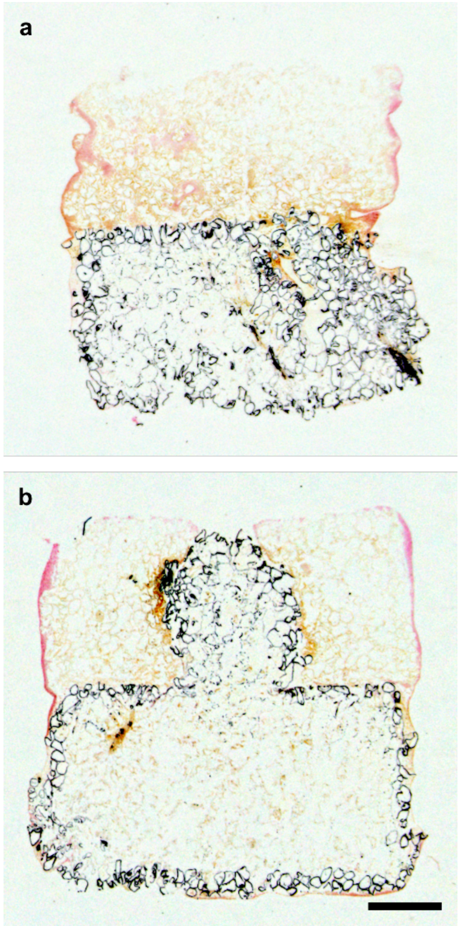


**Figure S4. Von Kossa staining of in vitro bone-connective tissue ETI. (a)** Cross-section of outer region of the interlocked composite. **(b)** Inner cross-section showing the peg and hole geometry of the stud and anti-stud subunits. Black = mineralized scaffold. Scale = 1 mm.

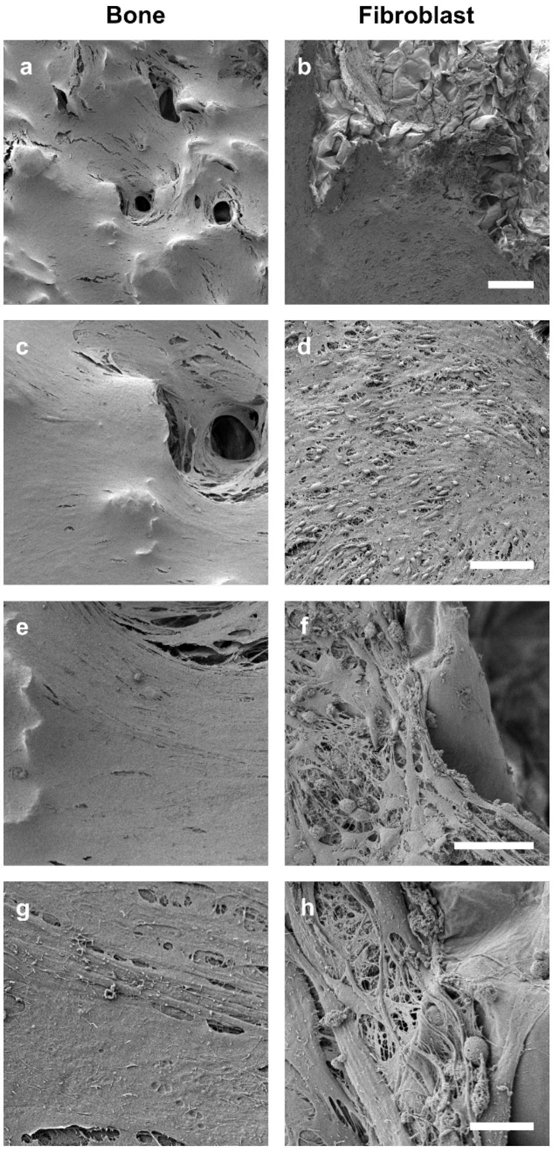


**Figure S5. SEM of bone-connective tissue ETI**. **(a,c,e,g)** MC 3T3 differentiated osteoblast seeded subunit. **(b,d,f,h)** NIH 3T3 fibroblast seeded subunit. Scale: **(a,b) =** 200 µm, **(c,d)** = 100 µm, **(e,f)** = 50 µm, **(g,h)** = 10 µm.

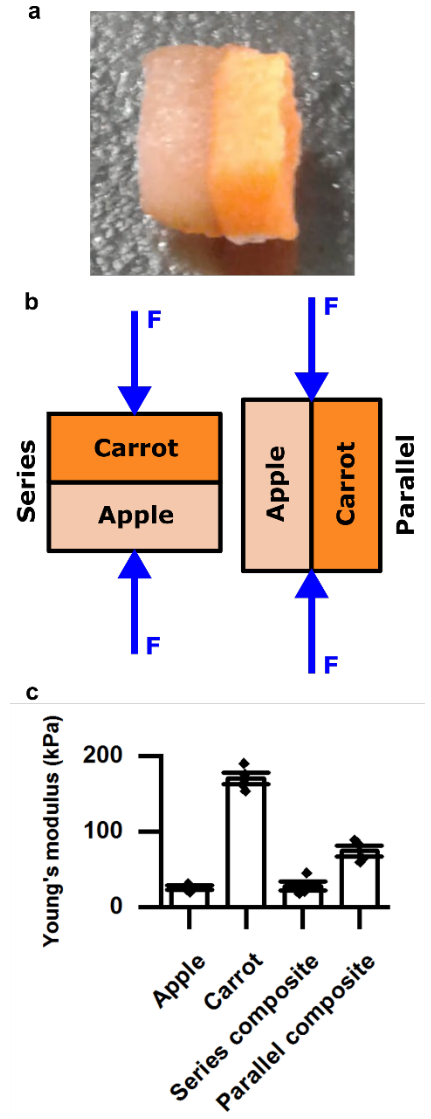


**Figure S6. Stress shielding with different source materials.** Apple and carrot-derived scaffolds were interlocked **(a)** and compressed in the series and parallel configurations **(b).** The effective Young's moduli for the materials and the isolated components **(c)**, N =4. The series and parallel composites were not statistically significantly different (P > 0.05) than the simplified model of two elastic rectangular prisms arranged in series $\left(\frac{1}{E_{mix}} = \frac{X_A}{E_A} + \frac{X_C}{E_C}\right)$ and parallel ($E_{mix} = X_A E_A + X_C E_C$) with individual moduli of the isolated apple and carrot components. All tissue interfaces in our bodies experience stress shielding when the mechanical environments of the constituents vary. The phenomenon of stress shielding is the difference in the stress applied to each body in a cohesive system.

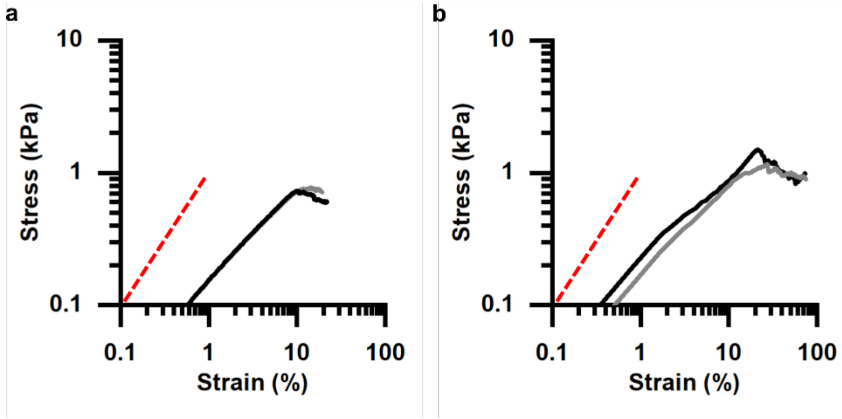


**Figure S7. Repeat controlled steady shear rate testing.** Controlled steady shear rate testing at a strain rate of **(a)** 0.001 $s^{-1}$ and **(b)** 0.01 $s^{-1}$. The black curve is the initial measurement, and the grey curve depicts a repeat trial on the same sample after a 5 min incubation period. The red dashed line is provided as a guide for the linear regime. The slow shear resulted in a failure point that was not as abrupt as it allowed time for more viscous dissipation. Reversible power-law behavior is observed at low strains: $\sigma \sim \varepsilon^y$ with (a) y = 0.72 ± 0.02 at 0.001 $s^{-1}$ and (b) y = 0.75 ± 0.02 at 0.01 $s^{-1}$.

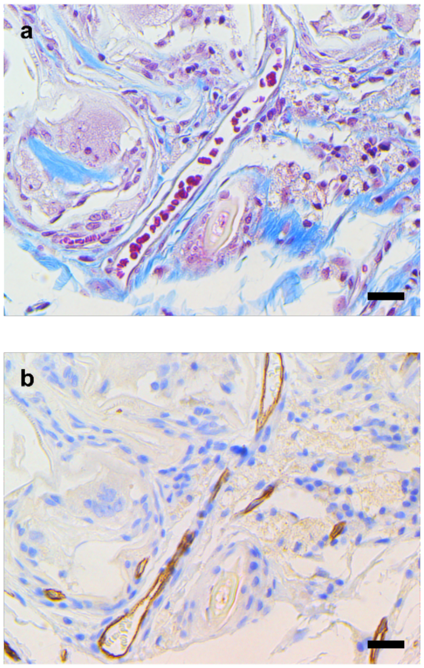


**Figure S8. Angiogenesis in the biomaterials in vivo. (a)** Masson's Trichrome staining showing red blood cells flowing through a large vessel inside the scaffold. **(b)** CD 31 staining for endothelial cells lining the blood vessel. Scale = 25 µm.

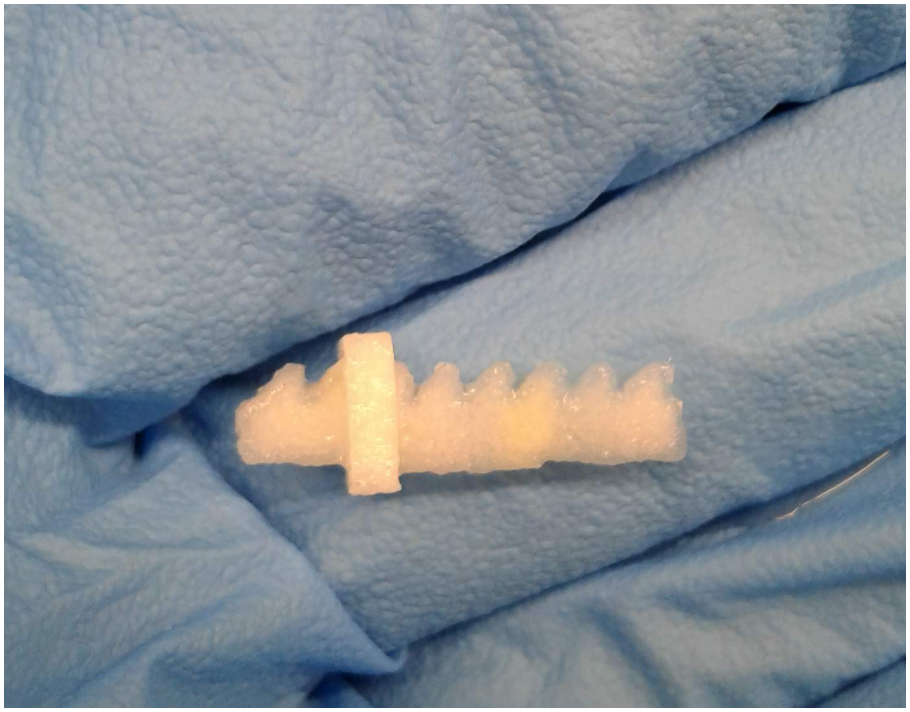

**Figure S9. Ratchet interlocking geometry.** A zip tie inspired ratchet strip and pawl were CNC milled out of apple hypanthium tissue. As highlighted here, a variety of interlocking geometries can be manufactured. The modular assembly of from custom subunits allows for different geometries and configurations to be created. Moreover, the use of building blocks can overcome significant challenges associated with using decellularized scaffolds, wherein the size limitations are a considerable scaling challenge. Modular assembly with controllable interfacial bioengineering can be used to circumvent the challenge of decellularizing large materials or sourcing large starting materials. The milled scaffold is displayed on a gloved hand for scale reference.

**Video S1. 3D milling with a computer numerical controlled (CNC) router.** A Shapeoko 3 CNC router with a 0.8 mm, 180° drill bit was used to carve McIntosh Red apples (Canada Fancy) into arrays of complementary stud and anti-stud geometries. Cutting was performed at a speed of 1 mm/s. The stud piece consisted of a 5 mm x 5 mm x 2mm base with a 2 mm cylindrical peg protruding from the center, and the anti-stud subunit had a 2 mm cylindrical hole in the center of the 5 mm x 5 mm x 2mm base. A Mandolin slicer was then used to cut the arrays of the subunits to their desired thickness.
